# Goggatomy: a Method for Opening Small Cuticular Compartments in Arthropods for Physiological Experiments

**DOI:** 10.1101/053298

**Authors:** Alan R. Kay, Davide Raccuglia, Jon Scholte, Elena Loukianova, Christopher Barwacz, Steven R. Armstrong, C. Allan Guymon, Michael N. Nitabach, Daniel F. Eberl

**Author notes:** Correspondence to Alan R. Kay.

## Abstract

Most sense organs of arthropods are ensconced in small exoskeletal compartments that hinder direct access to plasma membranes. We have developed a method for exposing live sensory and supporting cells in such structures. The technique uses a viscous light cured resin to embed and support the structure, which is then sliced with a sharp blade. We term the procedure a ‘goggatomy’, from the Khoisan word for a bug, *gogga*. To demonstrate the utility of the method we show that it can be used to expose the auditory chordotonal organs in the second antennal segment and the olfactory receptor neurons in the third antennal segment of *Drosophila melanogaster*, preserving the transduction machinery. The procedure can also be used on other small arthropods, like mites, *Daphnia*, mosquitoes, wasps and ants to expose a variety of cells.

## Introduction

Most arthropod sense organs are embedded in cuticular compartments and are relatively inaccessible. It is possible to record from bristle organ sensory cells after clipping off the bristle tip and applying an electrode to the cut end [1]. *Drosophila’s* Johnston’s organ (JO), which constitutes the fly’s ear is enclosed in a cuticular chamber, the second antennal segment (A2) and although it has been possible to record extracellularly from its afferent nerve process it has not as yet been possible to record intracellularly from the constituent cells of the chordotonal organ, as it has in larger insects [2, 3].

Although it is now possible to patch-clamp *Drosophila* CNS in semi-intact preparations [4], the exoskeleton has for the most part thwarted access to neurons and sensory structures buried within small cuticular compartments. Pioneering work by Dubin and Harris [5] showed that it was possible to obtain patch clamp recordings from olfactory sensory neurons in *Drosophila’s* third antennal segment, cut open with an iridectomy scissors. However, there is a need for a procedure that makes access simpler, more accurate and applicable to even smaller compartments. Providing open access to these compartments is essential if one is to be able to voltage clamp the cells, do single channel recordings or apply pharmacological agents.

In this communication we describe a method for opening the exoskeleton of arthropods that can be used to access small compartments not amenable to conventional microscopic dissection and expose, in a live state, the enclosed cells. There is a long history of using sliced arthropod eye preparations in neuroscience [6-8]. Our method modifies and extends this approach, by using a viscous photo- polymerizable resin that can be rapidly light cured, to support and anchor the cuticle. We believe that the method could be of value in a variety of physiological and pharmacological experiments on *Drosophila’s* JO and many other arthropod organs, appendages and brains.

## Materials and Methods

### *Drosophila* strains

For initial tests to develop the protocol, we used a Canton S wild type strain. To visualize membranes of all neurons in the fly, including those in the antenna, we used a *w elav^C155^-Gal4 UAS-mCD8-GFP* strain (Bloomington *Drosophila* Stock Center stock #5146). The *elav^C155^-Gal4* driver expresses in all neurons[9]. For Ca^2*^ imaging, we crossed *w elav^C155^-Gal4* females (Bloomington *Drosophila* Stock Center stock #458) with homozygous *w; UAS-GCaMP6m* males (Bloomington *Drosophila* Stock Center stock #42748). To record voltage-dependent fluorescence changes, we crossed homozygous *w; JO15-2-Gal4* females to *w; UAS-ArcLight^attP40^* homozygous males (Bloomington *Drosophila* Stock Center stock #51057)[10]. The *JO15-2* line, which expresses in the JO-A and JO-B subgroups of JO neurons, was derived from the original third chromosome *JO15* insertion [11, 12] by P-element remobilization to the second chromosome.

### *Drosophila* Saline

Our saline was based on the formulations of Wilson and Laurent (13) and Hardie et al. [14]. To optimize the recording we used a goggatomized *Drosophila* eye preparation where extracellular potentials in the retina were recorded with a glass electrode. Oxygenation of the saline proved unnecessary for sustaining the vitality of the preparations.

Drosophila saline (DS) in mM: 120 NaCl, 3 mM KCl, 1 CaCl_2_, 4 MgCl_2_, 4 NaHCO3, 1 NaH_2_PO_4_, 8 D-trehalose, 5 D-glucose, 2.5 L-alanine, 2.5 L-proline, 5 L-glutamine and 5 TES (pH 7.15)

### Imaging and Electrophysiology

Preparations were inserted into ~ 1mm ball of soft dental wax (white square ropes, Heraeu Kulzer, South Bend, IN) melted onto a 12 mm diameter cover glass which was placed in a perfusion chamber (Siskiyou Corp., Grants Pass, OR) and imaged on an Olympus BX50WI upright microscope equipped with a Hamamatsu ORCA-Flash 4.0 CMOS camera, with illumination provided by an X-Cite 120 LED (Excelitas Technologies Corp., Waltham, MA) through a Semrock (Rochester, NY) BrightLine filter set (472/30 Bandpass, 495 Dichroic and a 520/35 Bandpass) and controlled by MetaMorph software (Molecular Devices, Sunnyvale, CA).

The images in Fig. 9 were acquired on a Zeiss Axio Examiner upright microscope using a Plan Apochromat 40x N.A. 1.0 water immersion objective (Zeiss, Germany), using a Colibri LED system (Zeiss, Germany) with excitation at 470 nm. The objective C-mount image was projected onto the 80×80 pixel chip of a NeuroCCD-SM camera controlled by NeuroPlex software (RedShirtImaging, Decatur, GA). For image demagnification we used an Optem C-to-C mount 25-70-54 0.38x (Qioptiq LINOS, Fairport, NY).

Cells were stimulated with a glass microelectrode (~5M**Ω**) filled with DS which was controlled by pClamp software (version 9) through a Digidata 1322A (Molecular Devices, Sunnyvale, CA) coupled to an AMPI Iso-Flex stimulus isolator.

### LCR

The LCR composition: 70% BisGMA (bisphenol A diglycidyl methacrylate), 28.75% HEMA (2-hydroxy methacrylate), 1% EDMAB (2-ethyl dimethyl-4-aminobenzoate) and 0.25% CQ (camphorquinone).

LCR was cured with a SDI Radii Plus LED curing light with a light intensity of 1.5 W cm^−^2 and a peak at 460 nm.

A tungsten needle or electrode (~28 gauge and ~2” long) can be used for picking up a larger bead of LCR that can be used for the conventional goggatomy. For the free-arista-goggatomy smaller quantities of LCR are needed. In this case a tungsten needle can be coated with a thin layer of hard dental wax that the LCR wets (regular stick wax, Whip Mix Corp., Louisville, KY).

#### FTIR and DSC

Real Time FTIR studies were performed using Nicolet Fisher Nexus 670. Samples of the LCR were placed between two sodium chloride plates using 25 μm spacer beads. Conversion was evaluated using the absorption band at 1636 cm^−^1.

Differential Scanning Calorimetry studies were performed using a Perkin Elmer Diamond Differential Scanning Calorimeter modified to allow illumination of polymer samples. Heat evolved from the polymerization reaction was used to evaluate reaction behavior using a plain aluminum pan as a reference.

For both Real Time FTIR and DSC reactions were monitored for 3 minutes to evaluate reaction duration. All experiments were performed using a Rembrandt Allegro^TM^ lamp. A light intensity of 1.5 W cm^−^2 with peak irradiance at 450 nm.

All chemicals were from Sigma-Aldrich. All data is expresses as Mean ± SEM, with all the data measured from different preparations. All mean responses, unless otherwise noted, were significantly different from the baseline noise as judged by a two-tailed Student t test with p<0.001.

## Results

The method described here is simple; the body part is coated with a custom formulated light cured resin (LCR), which is applied as a viscous liquid and then cured to a solid by exposure to light. The sample embedded in the cured compound is then sliced with a fine razor blade while covered with a physiological saline (Fig. 1). The cured resin supports and reinforces the exoskeleton as the blade moves through it preventing its collapse and provides a handle for manipulating and positioning the sectioned material.

**Fig. 1.**
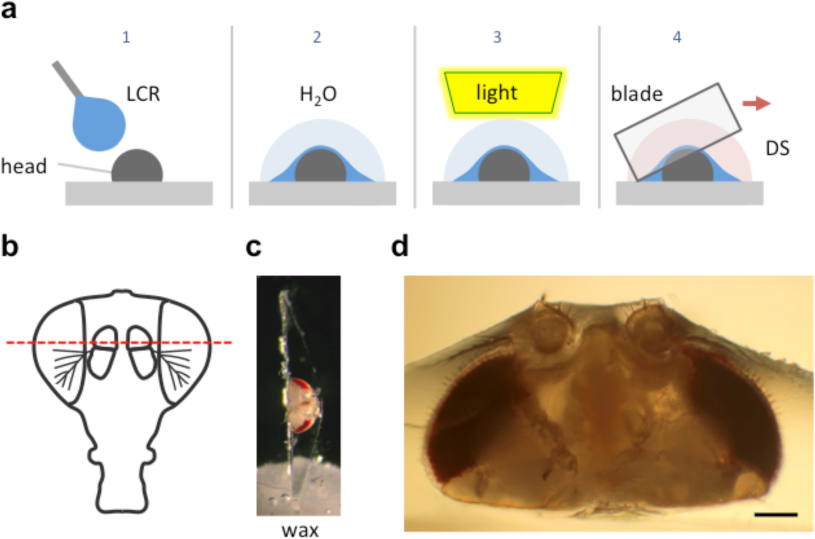
Schematic of the goggatomy procedure. (a) Sequence of steps illustrated for a *Drosophila* head (read left to right). The gray rectangle represents the surface of a plastic petri dish. (b) Approximate level of section to produce intact JOs. (c) Goggatomized head in chip of LCR inserted into wax. (d) Higher powered view of a different sectioned head. Scale bar 100 μm.

We have called the procedure a ‘goggatomy’, from the South African word for insect, gogga (*xoxo* pronounced ‘cho-cha’ where ‘ch’ is pronounced as in Scottish lo**ch**) which derives from the Khoisan language; the original inhabitants of South Africa who have a unique click based language [15].

The LCR used for this procedure is clear and becomes hard after a few seconds of irradiance with a blue light (460 nm). We will also show how it can be used to affix insects to a substrate and how it can be used to aid the viewing of neurons in intact animals.

We illustrate the method using the A2 of *Drosophila melanogaster*. The head of a cold-anesthetized fly is cut off with a blade and placed posterior side down on the cover of a 35 mm plastic petri dish. A small drop of LCR is applied to the anterior side of the head and allowed to flow over it. The droplet should be a little larger than the head (~2 μl) and is applied with a thin needle. Once the resin has covered the head a small drop of water (~10 μl) is placed over the LCR. The resin is then cured with a dental curing light held within a few millimeters of the LCR drop for 1 min. The water serves two purposes: (1) It reduces the amount of oxygen, which inhibits curing of the outer layer of resin. (2) It helps dissipate heat (*vide infra*). The water is wiped off the cured drop with a tissue and a drop of *Drosophila* saline (DS – see Methods) is placed over the cured LCR. A blade (Feather, carbon steel, Ted Pella Inc.), new and cleaned with ethanol, is broken into quarters and used to slice through the embedded head. Under a dissecting microscope, the blade is carefully oriented in the plane that one desires to cut. The blade is then firmly and quickly drawn through the encapsulated head, cleaving it into two pieces. As soon as the cuticle is breached, DS flows into the cut. The parts are securely embedded in the cured LCR and the separated LCR chips can be handled with forceps and placed in a dish with DS. The cured LCR is denser than water and sinks the sample to the bottom of the dish. To hold and position the sample we use a small piece of soft dental wax melted onto a 12 mm circular coverslip. An edge of the LCR chip containing the sample is simply pressed into the wax at the appropriate orientation.

We use the top of a 35 mm tissue culture dish as a work surface, since it forms an ideal platform that can be held and orientated with one hand while the other cuts through the embedded insect. Moreover the LCR bonds tightly to the surface of the dish so that the preparation remains fixed as the blade is drawn through it.

### LCR Composition and Properties

LCRs are widely used in dentistry to create permanent durable implants. The chemical components from which they are formulated have a long history of use in the human oral cavity and extensive tests have established their safety [16]. We screened five published formulations [17]. Of these, their resin #3 (see *Methods and Materials* for composition) proved best in terms of its ability to support the exoskeleton, retain the tissue after cutting and bond to the dish. The latter is important since the LCR should adhere to the dish as the blade moves through the sample. The cured resin fractures along the line of the cut rather than shattering. Moreover this formulation is optically clear with a refractive index of 1.57 that facilitates index-matching improving optical resolution. It is worth noting that light cured adhesives have been used to mount *Drosophila* for *in vivo* recordings [18, 19]

Polymerization of the LCR was followed by real time Fourier transform infrared spectroscopy. The average conversion of the resin was 68% (n=3) after 90 s of illumination. At this point the polymer matrix has vitrified so that the diffusion of residual monomer would be slow. Similar experiments performed with differential scanning calorimetry (DSC) found that most of the reaction occurred within the first 20s of illumination.

When the LCR polymerizes, its density increases and hence it shrinks. Under the conditions employed here, where an unconstrained thin layer is applied to the insect the shrinkage is likely to be nonuniform. The fact that it is cured under water might have an influence on this too. The surface of the cured LCR appears reticulated, which further supports a nonuniform curing process. Moreover after cutting through the A2, its profile does not appear to be deformed, as might occur if the shrinkage of the LCR squeezed the antenna appreciably.

Two important aspects of the LCR are its viscosity and its ability to wet arthropod cuticles. The high viscosity of the LCR (~1200 cP) allows one to suspend a droplet about the size of a fly’s head without it dropping off the applicator. The viscosity of the LCR slows its spread along the cuticle, which allows one to cover only part of the head or body, if so desired (*vide infra*). The wetting characteristics of the LCR induce it to penetrate between the setae and into even the finest crevices in the insect cuticle. This ensures that LCR holds the fly part, as the resin does not actually bond to the insect cuticle.

The LCR formulation used here does not generate much heat during the curing process. We tested the heat generated during curing by applying a drop of LCR to a piece of aluminum foil overlying a thermistor probe. Illuminating the LCR for 1 min led to an increase in temperature of only 2.6 ± 0.5 °C (n=4, starting temperature 24.3 ± 0.7 °C). For most physiological applications, this is a negligible increase.

### Morphology of Sensory Cells post-Goggatomy

To expose JOs, a fly head embedded in LCR was cut horizontally dorsal to the A2-A3 joint (Fig. 1b). At this level of section it was possible to observe intact scolopidia in the dorsal parts of both divided A2s. The form of the JO could be clearly resolved under bright field optics in goggatomized preparations, with the scolopidia attached to the remnants of the stalk (Fig. 2). The outer surface of the scolopale cells was bright and lustrous, suggesting that the scolopidia are preserved. Many scolopidia remain attached to the joint cuticle via their dendritic caps. Pushing against the scolopidium at right angles to its long axis with a patch electrode allows one to assess the integrity of this link. If the link is broken the scolopidium swings free.

**Fig. 2.**
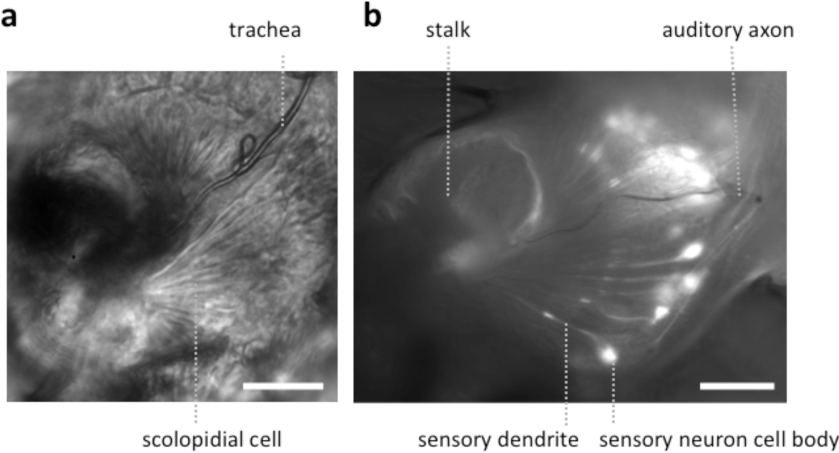
Light microscopic images of JOs within a goggatomized A2. (a) Bright field image of JO (b) Fluorescent image of JO expressing GCaMP6. Both specimens have approximately the same orientation. Scale bars 20 μm.

The goggatomy procedure allows preparations to be made that can be fixed, dehydrated, gold sputtered and viewed on a scanning electron microscope (Fig. 3). SEM images show that much of the scolopidial structure is preserved during goggatomy, even the dendritic caps connecting the scolopale cells to the stalk. The scolopidial cells are plump suggesting that the space is still filled. In some cases where the scolopidia have been cut, dendrites can be seen protruding.

**Fig. 3.**
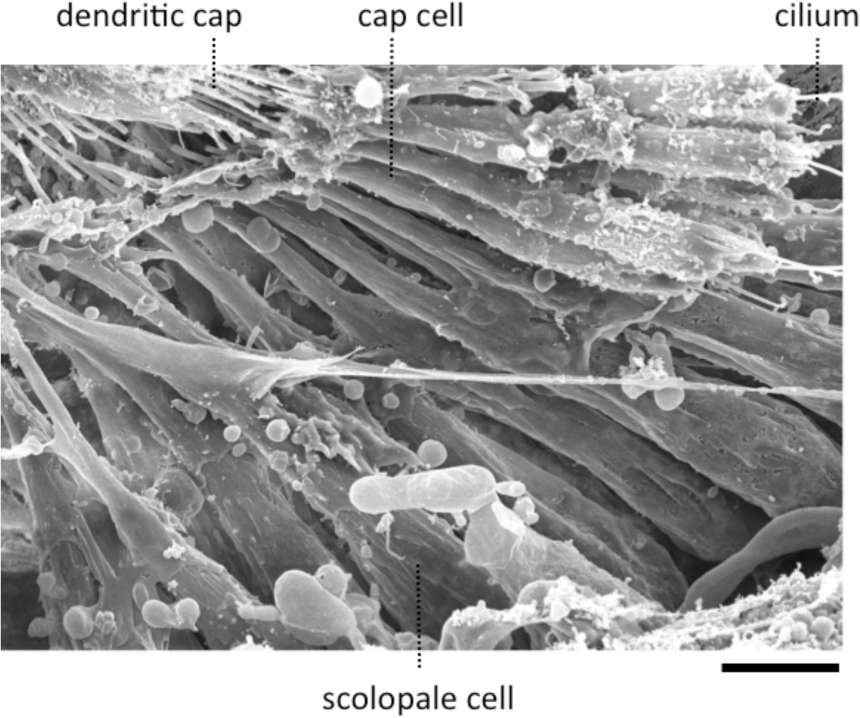
Scanning electron micrograph of a JO within a goggatomized A2. Scale bar 5 um.

The appearance of the scolopidia under SEM was not as good as that under bright field. We suspect that cells become fragile after dehydration and may fragment. Under saline in bright field microscopy, the full fan-like arrays of scolopidia can be seen (Fig. 2), whereas, under SEM this was less common.

### Physiology

In many cases when the head capsule is cut in the horizontal plane, the brain appears to pulse at a rate of ~ 1 Hz. This is due to the contraction of muscle 16, the frontal pulsatile organ [20, 21], whose pulsations can persist for up to 5 hours *in vitro*. Moreover, we have used fly lines that express GFP in neurons and in most cases the expression persists after many hours, again suggesting that the exposed cells are viable.

We have also assessed the toxicity of the procedure on whole flies. The dorsal surface of whole flies was attached to a plastic dish with LCR and the survival of the flies was monitored. Flies survived for more than 6 hours and remained fully motile to the extent that the tethering would allow. This survival resembles that when flies are held in pipette tips for extracellular electrophysiology [22]. This suggests that if any adhesive crosses the cuticle it has little toxic effect on the organism.

To assess the electrophysiological vitality of cells in the goggatomized preparations we used the fluorescent voltage sensor ArcLight developed by Pieribone and colleagues [23]. ArcLight is maximally fluorescent at hyperpolarized potentials, its fluorescence declining roughly linearly as the cell depolarizes. Moreover, its temporal response is sufficiently rapid to follow action potentials with reasonable fidelity. To determine if the sensory cells of JO had a hyperpolarized resting potential, characteristic of live cells, goggatomized A2s were perfused with a saline where all the Na* was substituted by K* (HiK), which should depolarize the cells to ~ 0 mV. Application of the saline led to a rapid decrease in fluorescence, consistent with depolarization (mean %ΔF/F = −28.5±3.2, n=16) (Fig. 4a). If the cells had no resting potential, application of HiK should lead to no or little change in the fluorescence of ArcLight. HiK stimulation induced similar changes in mushroom body neurons expressing ArcLight (mean %ΔF/F =-30.7 ± 5.1, n=12). Subjecting sensory neurons expressing GFP to HiK led to no significant change in fluorescence (mean %ΔF/F -0.02±1.4,n=9), indicating that that the changes in the ArcLight flies are not the result of the cells shrinking or swelling.

**Fig. 4.**
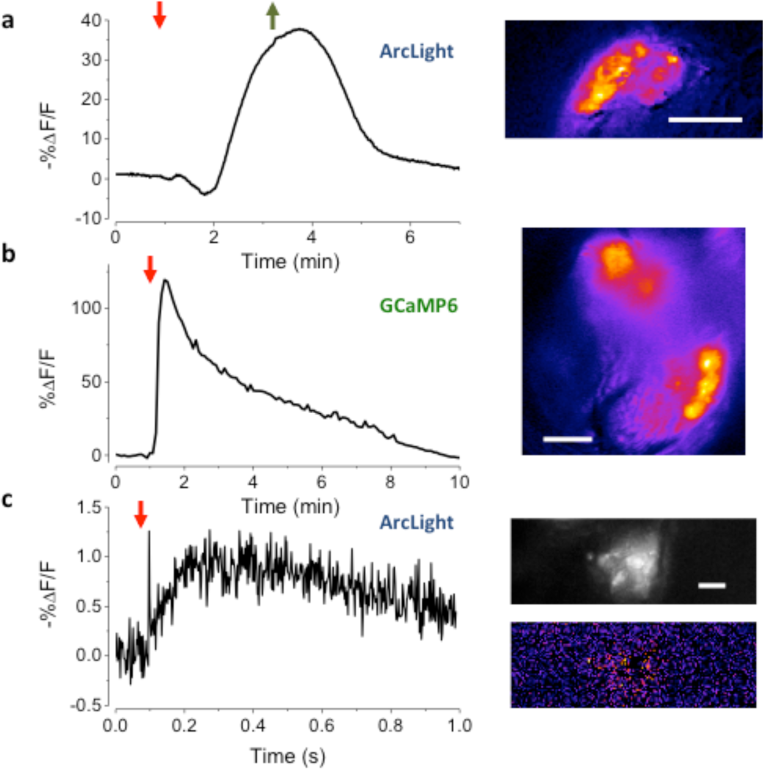
Assessing the vitality of exposed JOs. (a) Response of sensory cells in JO expressing ArcLight to the perfusion of HiK (red arrow) and restoration of normal saline (green arrow). Inset, pseudo-color image of difference between the image prior to stimulation and at its peak. Scale bar 20 μm. (b) Response of JO expressing GCaMP6 to application of pymetrozine (arrow, 3.4 μM). Inset, is pseudo-color difference image. Scale bar 20 μm. (c) Local electrical stimulation (arrow, 200 μs, 0.1 mA) of sensory cells in JO expressing ArcLight. Scale bar 10 μm.

To activate sensory neurons directly, a glass microelectrode was placed close to the cell bodies in a goggatomized A2s. When a pulse of current was delivered to the electrode, the fluorescence of the cell body declined and then increased back to baseline, consistent with depolarization and then repolarization of the cell (Fig. 4c, mean %ΔF/F= -1.5 ± 0.1, n=7).

We also used the insecticide pymetrozine to determine if the scolopidia survive the goggatomy procedure with their transduction mechanism intact. Pymetrozine interacts with the TRPV ion channel complex (with subunits Nan and Iav) resulting in a large influx of calcium [24]. Application of 15 μM of pymetrozine to goggatomized A2s from flies expressing GCaMP6 resulted in the consistent and rapid elevation of calcium in the sensory cells cells (Fig. 4b mean % ΔF/F = 56.3± 3.2 % ΔF/F, n=17). The response was irreversible and could not be restored after washing off the pymetrozine.

### Free-Arista Goggatomy

We modified the goggatomy procedure so as to expose the JO while preserving motion about the A2-A3 joint. The procedure is detailed in Fig. 5a and is termed the Free-Arista (FA) goggatomy. Directing a small short air pressure pulse towards the surface of the saline moved the arista. Movement of the arista was detected by making a high-speed video and activation of the JO by increases in calcium detected by GCaMP6 (Fig. 5b).

**Fig. 5.**
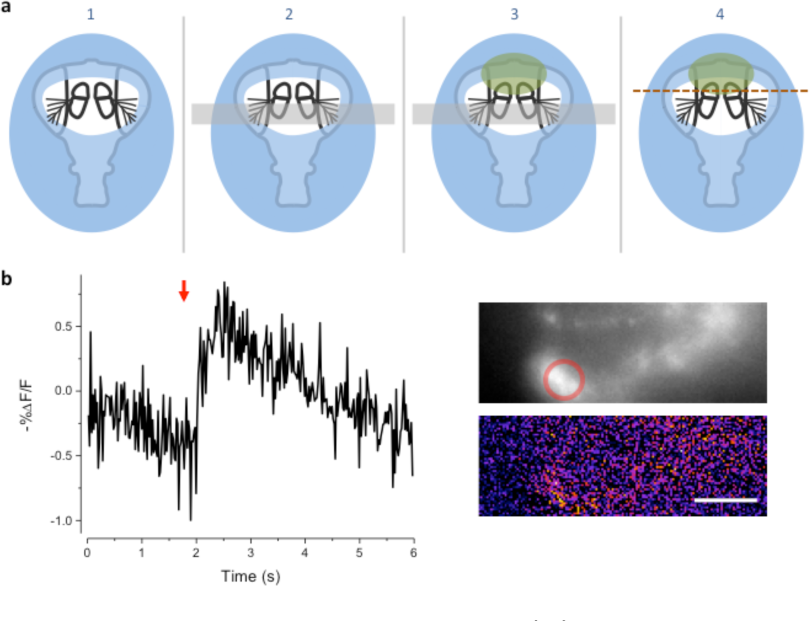
Free-Arista Goggatomy. (a) Schematic of the procedure (1) Place the cut head on a small droplet of LCR (blue) and allowed it to sink so that its edges, but not the antennae or arista, are covered. Cure for 20s. (2) Place a small strip of colored Cellophane (gray rectangle) over the A3, to prevent LCR from pulling the antennae up when it is applied to A2. (3) Apply a very small droplet of LCR so that it covers the dorsal margins of the A2 (green). Begin light curing, then remove the plastic strip and apply a droplet of water to the preparation. Cure for 1 min. Wick off the water with a tissue. (4) Apply DS and cut through the head as indicated by the dotted line. (b) Response of JO cells expressing GCaMP6 in a FA-goggatomized preparation to an air pulse (120 ms, ~ 1 psi) directed at the solution. Inset top, fluorescence, bottom pseudo-color difference image. Scale bar 20 μm.

### Olfactory receptors

We further modified the goggatomy procedure to section the A3 while preserving the olfactory receptor neurons (ORNs). The approach is detailed in Fig. 6a and shows how one can cut the antenna without embedding the sensory sensilla. Bath application of isoamyl-acetate led to a robust response in some of the sensory neurons, showing that their responsiveness is preserved Fig. 6c.

**Fig. 6.**
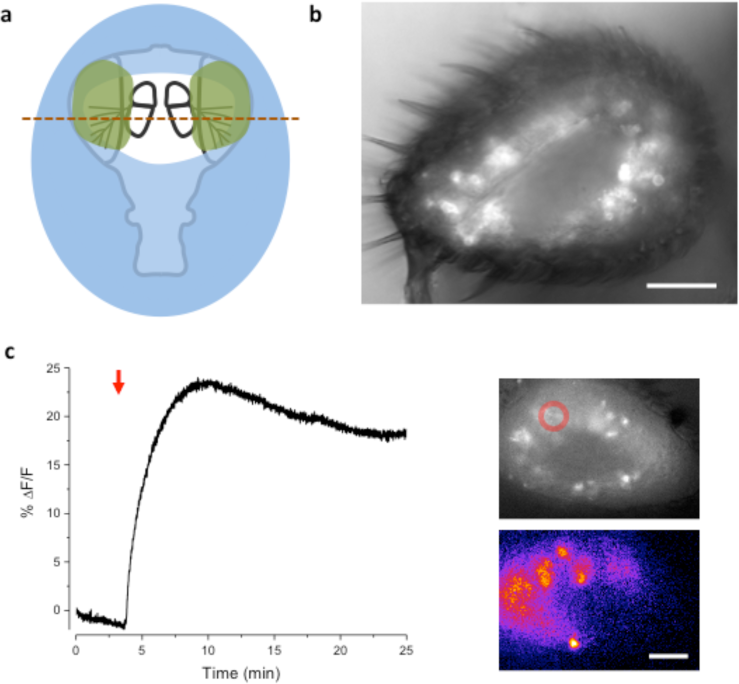
Sectioning A3 and response of ORNs to odorant. (a) Schematic of the procedure. Place the cut head on a droplet of LCR (blue) and allow sinking so that the margins of the head, but not the arista, become covered. Light cure for 20s. Apply small droplets of LCR over the arista and margins of the antennae (green). Note, the frontal surface of A3 should not be covered so that the sensory sensilla are free. Add droplet of water over preparation and cure for 1min. Wick off water with tissue. Add droplet of DS and cut horizontally through the A3s (red line). (b) Image of goggatomize A3 expressing GCaMP6. (c) Increase in intracellular calcium in ORNs expressing GCaMP6 to the application of iso-amyl acetate (arrow, a 5 μl drop of a 0.67 mM solution was added to the ~2 ml bath). Upper inset, fluorescence prior to odorant application. Lower inset pseudo-color difference image. Scale bar 20 μm.

### Air-Gogga Preparation

In some cases it might be desirable to produce a preparation where some part is not immersed in saline and exposed to the atmosphere. In the case of an antenna a goggatomized head prepared as in the last section is removed from the holding saline and placed on a piece of clear tape over a hole (~ 1 cm diameter) cut into a petri dish lid (Fig. 7). Vacuum grease is used to hold the section to the tape, exercising caution to avoid covering the antenna. The seal does not have to be complete as water tension prevents the saline from leaking out. The lid is inverted and a drop of saline is applied over the specimen. An opening is then cut in the tape overlying the nervous system with a blade tip allowing saline to flow in. We should note that we have not as yet tested whether, for example the A3 remains responsive to odorants.

**Fig. 7.**
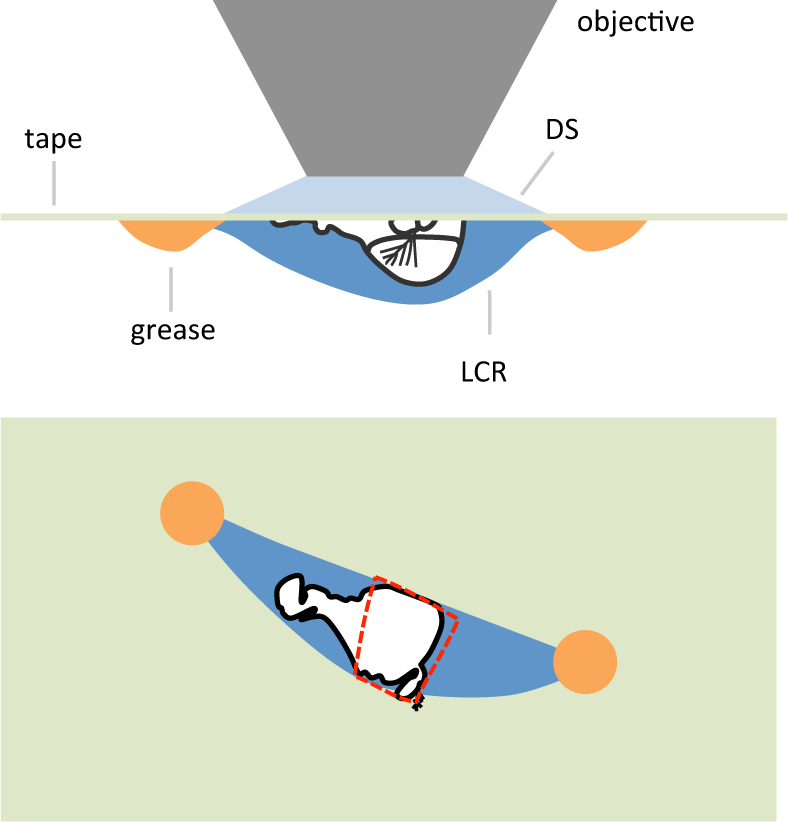
Schematic of an air-gogga preparation. Upper panel, side view. Lower panel, top down view. The dashed red line represents a hole cut into the tape to allow saline to enter the preparation.

### Other preparations

The goggatomy procedure is versatile enough that is can also be useful for a number of other exoskeletal compartments besides the antennae. As examples we show sensory cells in the proboscis and chordotonal organs in the legs (Fig. 8). Goggatomy also provides a convenient way of accessing the musculature and cells of the head, proboscis, halteres and sex organs.

**Fig. 8.**
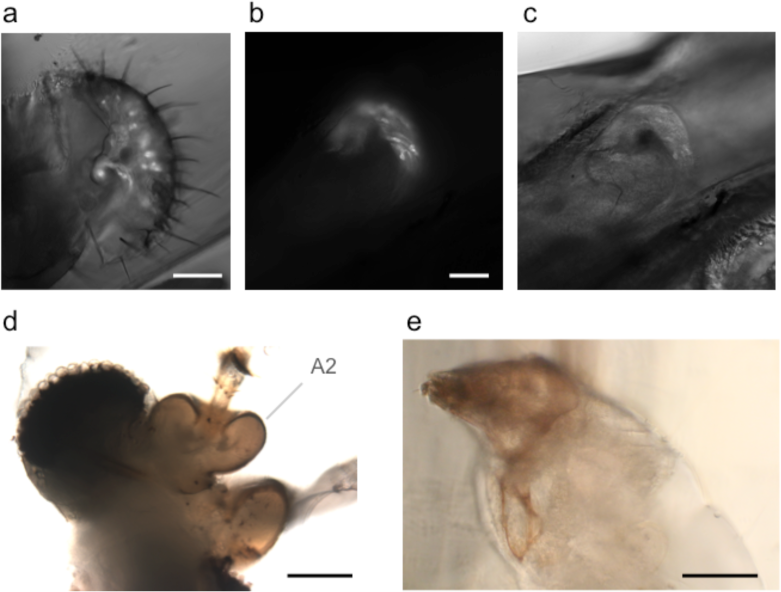
Goggatomy of Drosophila and other arthropods. (a) *Drosophila* labellum, GFP, scale bar 50 μm. (b&c) *Drosophila* leg chordotonal organ, GFP, scale bar 25 μm. (b) Fluorescence and (c) bright field. (d) Mosquito head, *Culex pipiens*, scale bar 50 μm. (e) Mimolette cheese mite, *Acarus siro*, scale bar 50 μm.

In addition, we have found that the goggatomy procedure is applicable to other small arthropods and have tried them on the following: ants, mosquitoes, daphnia and mites. Some examples are shown in Fig. 8.

### Using the LCR to view neurons in intact insects

We have found that we can use the LCR to mount whole live flies under a coverslip to view the antenna with a water immersion objective. The refractive index of cured LCR is 1.57; this makes it possible to preserve the high numerical aperture of the water immersion objectives. The procedure, which we term a ‘gogga cap’, is illustrated in (Fig. 9). A cold-anesthetized fly is affixed to a small piece of fishing line (0.25 mm diameter, ~ 4 mm length) that is held vertical on a small piece of poster tack under a dissection microscope. To do this, a drop of LCR is placed on the top of the fishing line and the ventral surface of the thorax is placed on the drop of resin. The fly is secured to the line by curing the LCR. A small drop of LCR is applied to a coverslip, which is attached to a coarse micromanipulator via a small rod. Under a stereomicroscope the cover slip is maneuvered over the head and lowered until the resin starts flowing over the head.

**Fig. 9.**
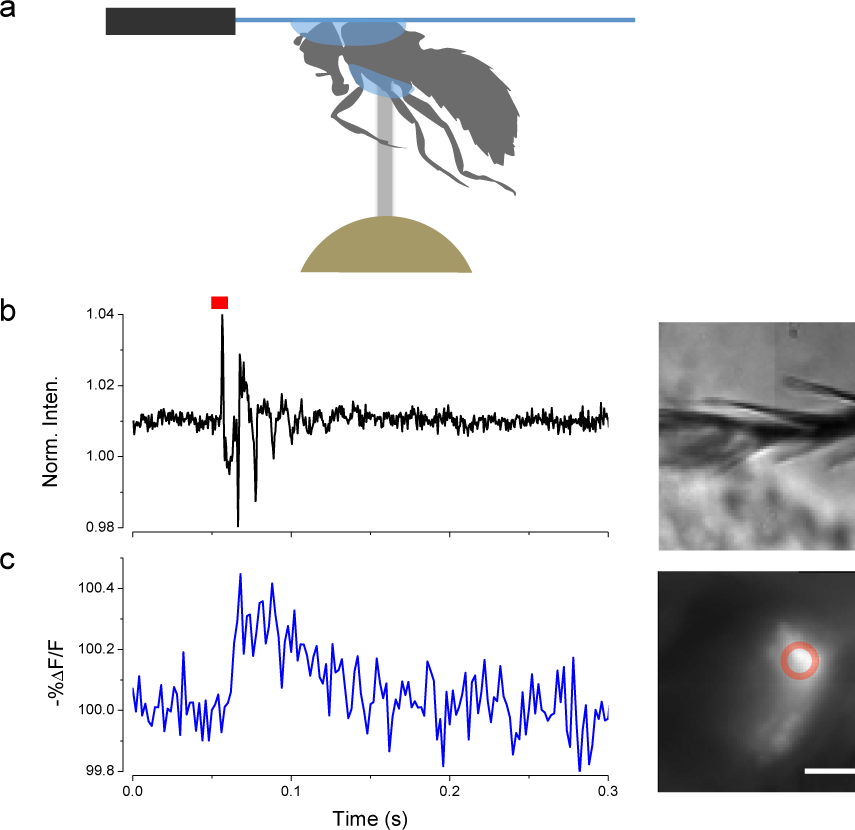
Mounting live flies with LCR. **(a)** Schematic of the mounting procedure. (b) Movement of arista (inset right) induced by a single sound pulse (red). (c) Response of JO expressing ArcLight in the same fly as (b). Scale bar 20 μm.

When the desired coverage is achieved the resin is cured. With this procedure it is possible to cover the 1^st^ and 2^nd^ antennal segments while leaving the 3^rd^ segment free.

In this configuration it is possible to image JO through the cuticle while stimulating the antenna with near field sound and detect changes in potential with ArcLight. An example of such a recording is shown in (Fig. 9).

## Discussion

We have developed a simple procedure for opening up the exoskeleton of arthropods, which exposes the live tissue and neuronal components. The goggatomy procedure opens up previously inaccessible cells for direct physiological and pharmacological manipulation.

We have used the fluorescent voltage sensor ArcLight to show that sensory neurons in JO have a substantial negative potential and auditory sensory neurons respond to current injection. The sensory transduction apparatus of the both JO and ORNs remains intact as judged by chemical stimulation. Moreover, in a preparation where the A2-A3 joint is preserved, sensory neurons respond to joint displacement. In addition, we have found that in *Drosophila* brain exposed by goggatomy, spontaneous and rhythmic activity persists *in vitro* [25].

For arthropods where it is not feasible to use genetically encoded indicators, the goggatomy procedure makes it possible to use the acetoxylmethyl ester (AM) loading of synthetic ion-indicators like fluo-4 into cells [26] as has been done in the honeybee [27]. *Drosophila* neurons can load with calcein-AM [28], however some synthetic calcium probes may not load very efficiently (Jing Wang, personal communication).

The method does not rely on the LCR adhering to the cuticle. Arthropod cuticles have a thin wax layer and the resin in the uncured form wets it, but when cured does not bond to it. The specimen is held in place because the resin forms a replica of the exoskeleton surrounding hairs and filling tiny gaps. The method is not suitable for soft-bodied animals, like *C elegans* or *Drosophila* larvae, since once sliced the specimen falls out of the resin. It is worth noting that the goggatomy procedure can facilitate the immunocytochemical staining of small arthropods since it makes the sections dense and provides a handle for manipulating the tissue.

Goggatomization of *Drosophila* heads exposes the beating frontal pulsatile organ (muscle 16) that can be sustained *in vitro* for a few hours. We suggest that this preparation could be a useful one for studying the rhythmicity of the heart. Moreover, the preparation is simple enough to be used in classroom demonstrations and student labs.

More than ten years ago Wilson, Turner and Laurent [4] showed how it was possible to perform patch clamp recordings on adult *Drosophila* brains. Their method has been widely used opening up this important model organism to physiological investigation. The first author spent two years trying to get intracellular recordings from scolopidia without success. The sensory neuron cell bodies are covered by a tough extracellular matrix, which hinders both patch and sharp recordings. The scolopale cells are more rigid than other cells, thwarting attempts to patch and penetrate the cells. Although we were unable to make direct intracellular recordings from scolopidia, we believe that it would be imprudent to suggest that it is impossible. It might be possible to make intracellular recording from the components cells of scolopidia with an appropriate sharp electrode, perhaps quartz or by digesting the extracellular matrix with proteases. Our confidence that intracellular recordings can be made from *Drosophila* JO is bolstered by reports of wholeMcell and perforated patch recordings from ORNs in A3s that have been cut open [5, 29]

Isolated neuronal preparations have played an important role in the progress of neuroscience. These include among others, the squid giant axon and synapse, the limulus eye, and isolated gastropod ganglia. We believe that the goggatomy procedure will be of great value in helping to reveal the secrets of sensory organs and other cells trapped with the confines of very small cuticular compartments.

## Appendix

## Goggatomy - step by step

1. Clean blade with alcohol and snap into quarters.
2. Melt a small piece of soft dental wax onto a 12 mm circular coverslip, place in a 35mm petri dish filled with DS (Fig. A 1 a).
3. Cold anesthetize flies for less than 30s
4. Pick up a drop of LCR (volume just a little larger than the head) with a tungsten tip.
5. Lie fly on its side on the lid of the 35 mm petri dish. Cut off head with blade. Place head on its posterior surface.
6. Apply a droplet of LCR on the head to completely cover it (Fig. A 1 b).
7. Place a drop of water on the LCR and apply the curing light for 1min, protecting eyes with goggles (Fig. A 1c).
8. Wipe off the saline with a tissue and apply a drop of DS.
9. Under higher magnification orient the fixed fly head. Hold the blade at an angle of ~ 30^o^ to petri dish surface (Fig. A 1d). Cut through the embedded head just above the A2-A3 (Fig. 1b) joint with a swift and firm motion.
10. Trim the cured LCR chip (Fig. A 1e) and transfer to the holding dish.
11. Insert the chip into the dental wax under saline, with the cut surface appropriately oriented (Fig. A 1f).

**Fig. A 1.**
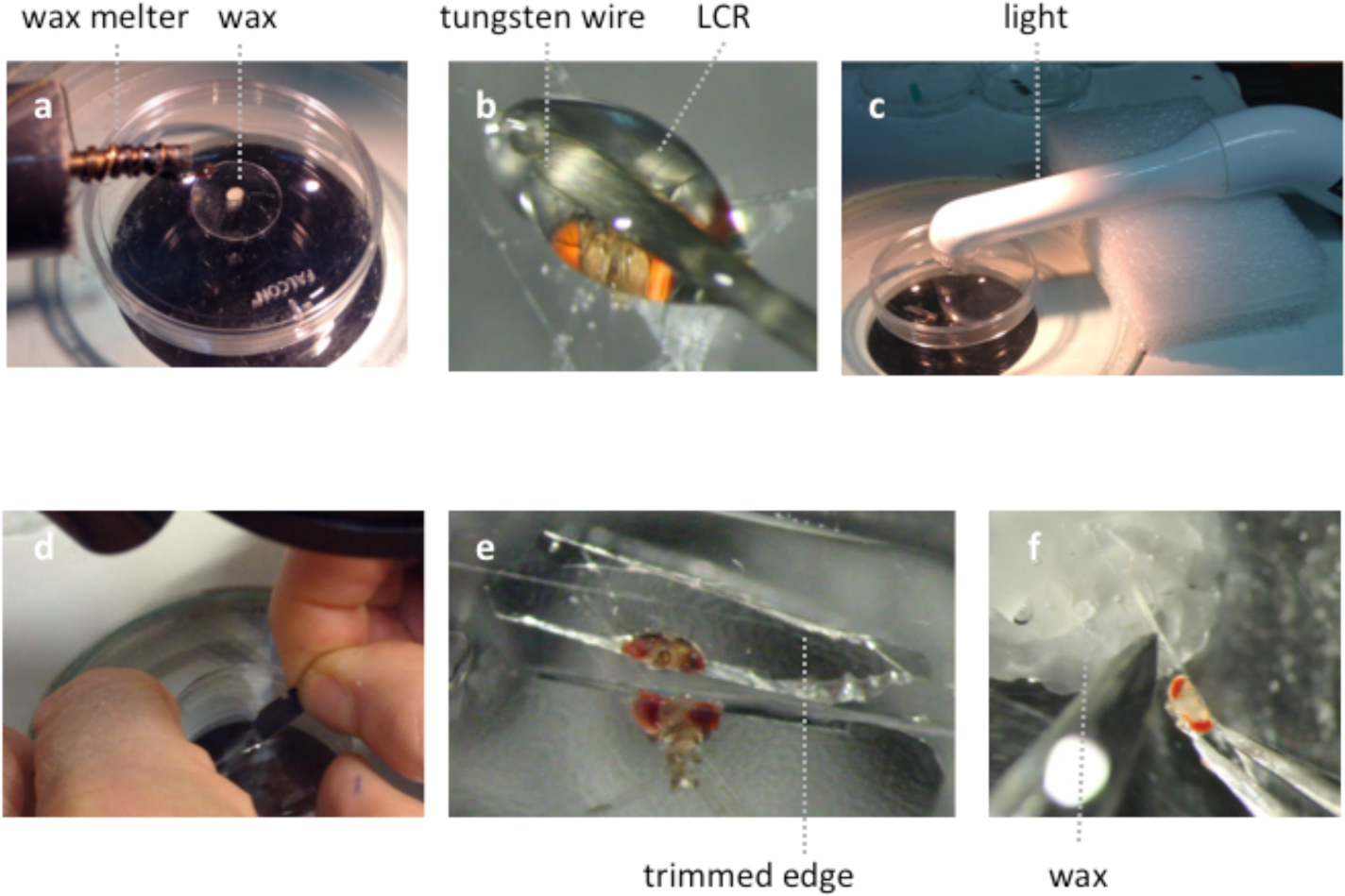
Illustration of the goggatomy procedure on a *Drosophila* head. (a) Melting a small piece of soft dental wax onto a circular cover slip. (b) Applying LCR to *Drosophila* head. (c) Curing light is place close to specimen. (d) Cutting through cured LCR. (e) Trimming embedded sections. (f) Placing chip in wax.

## Author Contributions

ARK & DFE initiated the project. ARK performed the experiments. ARK, DR & DFE analyzed the data. DFE, DR & MNN produced the fly lines. ES performed the SEM. JS, CAG & SRA made and analyzed the LCR. MNN, CAB, JS, CAG & SRA provided reagents. ARK wrote the paper with contributions from all other authors.

## Acknowledgments

We thank Mark Reagan and Matthew Wortel for measuring the refractive index of the LCR, Lyric Bartholomay for mosquitoes, Andrew Forbes for wasps and the New Pioneer Coop for the mimolette cheese, and Roger Hardie, Toshinori Kitamoto and Jing Wang for comments on an earlier version of the manuscript.

The work was in part supported by an award from the Office of the Vice President for Research, University of Iowa (ARK).

## References

1. Corfas G, Dudai Y. Adaptation and fatigue of a mechanosensory neuron in wild-type Drosophila and in memory mutants. J Neurosci. 1990;10(2):491–9. PubMed PMID: 2154560.

2. Field LH, Matheson T. Chordotonal organs of insects. Advances in Insect Physiology. 1998:1–230.

3. Hill KG. The physiology of locust auditory receptors I. Discrete depolarizations of receptor cells. J Comp Physiol. 1983;152:475–82.

4. Wilson RI, Turner GC, Laurent G. Transformation of olfactory representations in the Drosophila antennal lobe. Science. 2004;303(5656):366–70. doi: 10.1126/science.1090782. PubMed PMID: 14684826.

5. Dubin AE, Harris GL. Voltage-activated and odor-modulated conductances in olfactory neurons of Drosophila melanogaster. J Neurobiol. 1997;32(1):123–37. PubMed PMID: 8989668.

6. Hartline HK, Wagner HG, Macnichol EF, Jr. The peripheral origin of nervous activity in the visual system. Cold Spring Harb Symp Quant Biol. 1952;17:125–41. PubMed PMID: 13049160.

7. Hadjilazaro B, Baumann F. Afterpotentials of the visual cell of the honey-bee drone. Helv Physiol Pharmacol Acta. 1968;26(3):CR351–2. PubMed PMID: 5719361.

8. Wu CF, Pak WL. Quantal basis of photoreceptor spectral sensitivity of Drosophila melanogaster. J Gen Physiol. 1975;66(2):149–68. PubMed PMID: 809537; PubMed Central PMCID: PMCPMC2226201.

9. Lin DM, Goodman CS. Ectopic and increased expression of Fasciclin II alters motoneuron growth cone guidance. Neuron. 1994;13(3):507–23. PubMed PMID: 7917288.

10. Cao G, Platisa J, Pieribone VA, Raccuglia D, Kunst M, Nitabach MN. Genetically Targeted Optical Electrophysiology in Intact Neural Circuits. Cell. 2013;154(4):904–13. doi: Doi 10.1016/J.Cell.2013.07.027. PubMed PMID: WOS:000323202500019.

11. Sharma Y, Cheung U, Larsen EW, Eberl DF. PPTGAL, a convenient Gal4 P-element vector for testing expression of enhancer fragments in drosophila. Genesis. 2002;34(1-2):115–8. doi: 10.1002/gene.10127. PubMed PMID: 12324963; PubMed Central PMCID: PMCPMC1805626.

12. Kamikouchi A, Shimada T, Ito K. Comprehensive classification of the auditory sensory projections in the brain of the fruit fly Drosophila melanogaster. J Comp Neurol. 2006;499(3):317–56. doi: 10.1002/cne.21075. PubMed PMID: 16998934.

13. Wilson RI, Laurent G. Role of GABAergic inhibition in shaping odor-evoked spatiotemporal patterns in the Drosophila antennal lobe. J Neurosci. 2005;25(40):9069–79. doi: 10.1523/JNEUROSCI.2070-05.2005. PubMed PMID: 16207866.

14. Hardie RC, Martin F, Cochrane GW, Juusola M, Georgiev P, Raghu P. Molecular basis of amplification in Drosophila phototransduction: roles for G protein, phospholipase C, and diacylglycerol kinase. Neuron. 2002;36(4):689–701. PubMed PMID: 12441057.

15. Haacke WHG, Eiseb E. A Khoekhoegowab Dictionary. Windhoek, Namibia.: Gamsberg MacMillan Pub.; 2002. 740 p.

16. Pereira SA, de Menezes FC, Rocha- Rodrigues DB, Alves JB. Pulp reactions in human teeth capped with self-etching or total-etching adhesive systems. Quintessence Int. 2009;40(6):491–6. PubMed PMID: 19587890.

17. Pashley DH, Tay FR, Carvalho RM, Rueggeberg FA, Agee KA, Carrilho M, et al. From dry bonding to water-wet bonding to ethanol-wet bonding. A review of the interactions between dentin matrix and solvated resins using a macromodel of the hybrid layer. Am J Dent. 2007;20(1):7–20. PubMed PMID: 17380802.

18. Seelig JD, Chiappe ME, Lott GK, Dutta A, Osborne JE, Reiser MB, et al. Two-photon calcium imaging from head-fixed Drosophila during optomotor walking behavior. Nat Methods. 2010;7(7):535–40. doi: 10.1038/nmeth.1468. PubMed PMID: 20526346; PubMed Central PMCID: PMCPMC2945246.

19. Budick SA, Reiser MB, Dickinson MH. The role of visual and mechanosensory cues in structuring forward flight in Drosophila melanogaster. J Exp Biol. 2007;210(Pt 23):4092–103. doi: 10.1242/jeb.006502. PubMed PMID: 18025010.

20. Murthy M, Turner G. In vivo whole-cell recordings in the Drosophila brain. In: Zhang B, Freeman MR, Waddell S, editors. Drosophila Neurobiology: A laboratory manual. New York: Cold Spring Harbor Laboratory Press; 2010. p. 534.

21. Demerec M. Biology of Drosophila. New York, NY: Hafner Publishing Co. Inc; 1950.

22. Eberl DF, Hardy RW, Kernan MJ. Genetically similar transduction mechanisms for touch and hearing in Drosophila. J Neurosci. 2000;20(16):5981–8. PubMed PMID: 10934246.

23. Jin L, Han Z, Platisa J, Wooltorton JRA, Cohen LB, Pieribone VA. Single action potentials and subthreshold electrical events imaged in neurons with a fluorescent protein voltage probe. Neuron. 2012;75(5):779–85. doi: 10.1016/j.neuron.2012.06.040. PubMed PMID: 22958819.

24. Nesterov A, Spalthoff C, Kandasamy R, Katana R, Rankl NB, Andres M, et al. TRP Channels in Insect Stretch Receptors as Insecticide Targets. Neuron. 2015;86(3):665–71. doi: 10.1016/j.neuron.2015.04.001. PubMed PMID: 25950634.

25. Rosay P, Armstrong JD, Wang Z, Kaiser K. Synchronized neural activity in the Drosophila memory centers and its modulation by amnesiac. Neuron. 2001;30(3):759–70. PubMed PMID: 11430809.

26. Grienberger C, Konnerth A. Imaging calcium in neurons. Neuron. 2012;73(5):862–85. doi: 10.1016/j.neuron.2012.02.011. PubMed PMID: 22405199.

27. Galizia CG, Sachse S, Rappert A, Menzel R. The glomerular code for odor representation is species specific in the honeybee Apis mellifera. Nat Neurosci. 1999;2(5):473–8. doi: 10.1038/8144. PubMed PMID: 10321253.

28. Li C, Meinertzhagen IA. Conditions for the primary culture of eye imaginal discs from Drosophila melanogaster. J Neurobiol. 1995;28(3):363–80. doi: 10.1002/neu.480280309. PubMed PMID: 8568517.

29. Cao LH, Jing BY, Yang D, Zeng X, Shen Y, Tu Y, et al. Distinct signaling of Drosophila chemoreceptors in olfactory sensory neurons. Proc Natl Acad Sci U S A. 2016;113(7):E902–11. doi: 10.1073/pnas.1518329113. PubMed PMID: 26831094.

